# Mammogram Segmentation using Multi-atlas Deformable Registration

**DOI:** 10.1101/542217

**Authors:** Manish Kumar Sharma, Mainak Jas, Vikrant Karale, Anup Sadhu, Sudipta Mukhopadhyay

## Abstract

Accurate breast region segmentation is an important step in various automated algorithms involving detection of lesions like masses and microcalcifications, and efficient telemammography. Existing segmentation algorithms underperform due to variations in image quality and shape of the breast region. In this paper, we propose to segment breast region by combining data-driven clustering with deformable image registration. In the first phase of the approach, we identify atlas images from a dataset of mammograms using data-driven clustering. Then, we segment these atlas images and use in the next phase of the algorithm. The second phase is atlas-based registration. For a candidate image, we find the most similar atlas image from the set of atlases identified in phase one. We deform the selected atlas image to match the given test image using the Demon’s registration algorithm. Then, the segmentation mask of the deformed atlas is transferred to the mammogram in consideration. Finally, we refine the segmentation mask with some morphological operations in order to obtain accurate breast region boundary. We evaluated the performance of our method using ground-truth segmentation masks verified by an expert radiologist. We compared the proposed method with three existing state-of-the-art algorithms for breast region segmentation and the proposed approach outperformed all three in most of the cases.

## I. INTRODUCTION

Automatic segmentation of Breast Region is a critical step in Computer Aided Diagnostic (CAD) tools for mammograms - with applications including detection of lesions, classification of lesions and diagnosis of breast cancer. It enables algorithms downstream in the CAD pipeline to be more efficient and accurate by constraining the analysis to be confined to a region of interest relevant to the algorithm.

Early attempts at breast region segmentation were based on thresholding pixel intensity histograms [1–5]. These methods estimate the intensity level which best separates the background from the foreground. Although these algorithms are computationally efficient, medical images are rarely strictly bimodal thus violating the fundamental assumption of these algorithms. Furthermore, thresholding algorithms do not take into account the relationship amongst pixels. Region growing algorithms [5, 6] overcome this limitation by assuming that the region to be segmented is connected. For example, Dehghani *et al*. [5] proposed a combination of thresholding and region growing operation. More recently, pattern analysis-based techniques [7–9] have attempted to automatically classify pixels as either belonging to the foreground or the background. The watershed algorithm [10, 11] is another approach where the image is treated as a 3D topography (with intensity level as height). This topography is flooded and the watershed lines which separate neighboring basins are used to segment the image. Deformable methods comprise another category of segmentation algorithms for the breast region. These offer finer accuracy by taking into account the shape of the segmented region by deforming contours or objects while minimizing an energy functional. Atlas-based segmentation, active contours [12], snakes [13], active shapes [14, 15] and level sets [16–18] are examples of this approach. A brief overview of the recent advancements in segmentation of the breast boundary is provided by Mustra *et al*. [19].

Despite the fact that image segmentation is a well-studied problem for several years, breast region segmentation still poses a challenge due to technical and physiological reasons. In this article, we will consider atlas-based image segmentation - a technique successfully applied in several imaging domains and modalities - neuroimaging [20], cardiac images [21], pulmonary imaging [22] *etc*. However, this has not yet been widely explored for breast region segmentation in mammograms. Jas *et al*. [23], in a previous conference paper, proposed an atlas-based breast region segmentation where landmark-based rigid registration followed by deformable image registration was applied. However, such a method crucially depends on the accuracy of the landmarks detected and the initial distance between the two images used for registration. In the case of mammography, automated landmark detection algorithms can detect at best a few points: the nipple [24], pectoral muscle edge [25], etc. Thus, a fully automated method is not feasible. Particularly because the breast tissue type varies considerably from individual to individual. The mini-MIAS [26] database, for instance, classifies the images into dense, glandular or fatty images. Thus, a strategy based on registration with a single atlas will most likely fail in this case. The tissue type (dense, glandular, or fatty) which predominately constitutes the breast region may affect the contrast of the image. The shape and size of the breast region also vary from individual to individual. A strategy based on multiple atlases instead of a single atlas will be more successful in such a scenario. Therefore, in this paper a multi-atlas based approach has been proposed for breast region segmentation.

**Figure 1:**
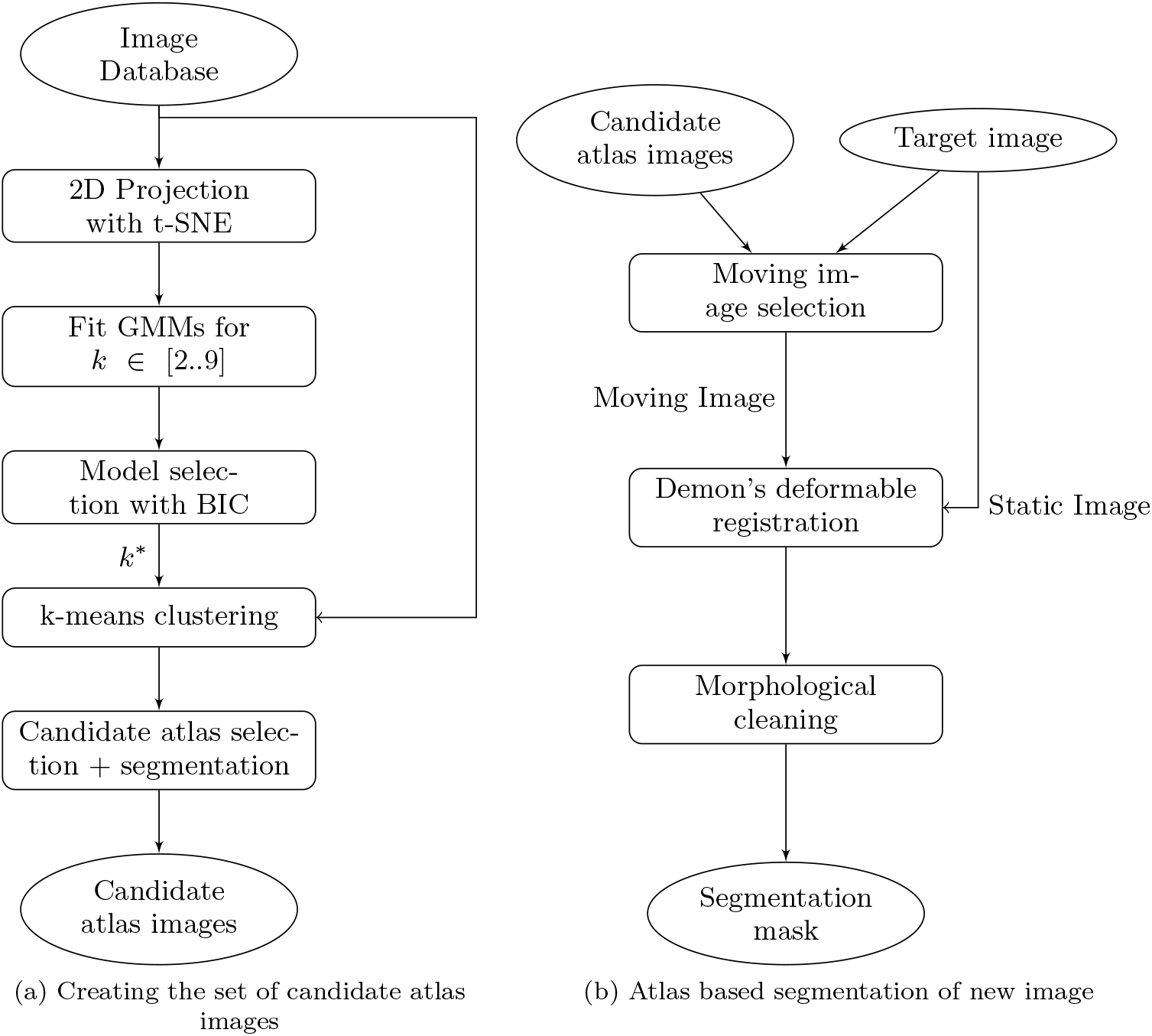
Schematic owchart for atlas-based segmentation with automatic atlas selection

In a multi-atlas strategy, one could either fuse segmentations obtained with different atlas images or generate one segmentation corresponding to the best atlas. Indeed, the selection of an optimal atlas has been demonstrated to affect the final segmentation accuracy by Wu *et al*. [27]. In this paper, we focus on the latter approach. We demonstrate that generating a database of atlases in a multi-atlas framework helps generate better segmentation masks. We formulate an automated breast region segmentation framework which uses two steps: first, selection of atlas and second, registration process and segmentation. We propose to create a data-driven library of atlas images. This is done by first identifying the types of images in the database and then selecting an optimal number of atlas images based on this information. Given a new image, its nearest matching atlas image is found from the set of candidate atlases and used for registration. The computed segmentation results are evaluated against ground-truth segmentation masks verified by an expert radiologist. We compare this approach against three competing methods on standard evaluation metrics and find that the proposed approach is able to outperform these by a large margin.

## II. METHODS

The proposed method involves two steps: first, selection of atlas and second, registration process and segmentation.

### A. Atlas Selection

The atlas selection procedure is data-driven. The efficiency and accuracy of the registration process depends on the atlas image used. Hence, instead of selecting atlas images randomly, we use clustering to determine which images will be the most representative ones. To efficiently compute the clusters as well as for visualization, we start by projecting the high-dimensional images to a 2D plane using t-distributed Stochastic Neighbor Embedding (t-SNE) [28]. The t-SNE method claims to preserve the inter-class separability while projecting to lower dimensions. As in the original work, we first reduce the dimensionality to 25 using Principal Component Analysis (PCA) to reduce noise and to compute the projection efficiently.

This data was then projected to a 2D space using t-SNE. We fitted Gaussian Mixture Models (GMM) with *k* Gaussians, *k* ∈ [2..9], to this 2D projected data. The optimal value of *k* will give the optimal number of clusters *k*\*. Bayesian Information Criteria (BIC) [29] is a popularly used technique in model selection. Given a set of observations with sample size *n*, each GMM has a maximized value of likelihood function, 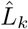. The BIC criteria in Equation (1) uses this value and the dimensionality of the GMM (the number of Gaussians), *k*, to find the most optimal model. The dimensionality, *k*, is used as a penalty term so that the model with highest dimensions is not always preferred. The model with minimum value of BIC gives the optimal number of clusters, *k*\*, Equation (2).
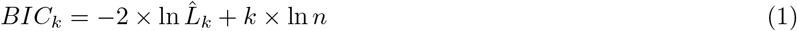

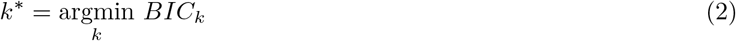

We used the value of *k*\* obtained above to perform a k-means clustering on the actual images. The next step is to identify an atlas image from each cluster. Since the central cluster image is the average of all images belonging to that cluster, it is blurred thus and not suitable for registration. We therefore picked the image closest to the cluster center as the best representative atlas image for that cluster. The image closest to the cluster center *C_k_* is the one which has the lowest Euclidean distance from *C_k_*. Given images {*I*_*k*1_, *I*_*k*2_…} belonging to the *k*th cluster, each with dimensions *W* × *H*, the corresponding candidate atlas, *I_atlas, k_*, is selected as given by Equation (3).
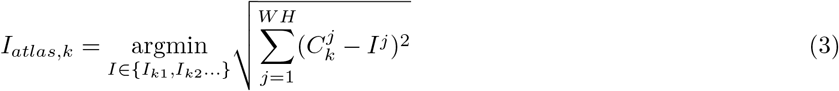

The candidate atlas images should be accompanied by ground-truth masks just as in any atlas-based segmentation procedure. This step is typically manual even though the segmentation procedure is entirely automatic. For the convenience of the reader, the atlas selection procedure is summarized in Figure 1a.

### B. Demon’s Registration

These candidate atlases are now used for the registration and segmentation. We used Demon’s Deformable registration [30] for the registration process. The complete procedure for the registration is shown in Figure 1b. Demon’s registration requires two images: a static image, and a moving image. The moving image is transformed to match the static image during the registration procedure. As opposed to affine registration where the same transformation is applied on the entire image, in deformable registration, each pixel in the image can be transformed independently. Given a new image for segmentation, we consider it as the static image, *S*, and need to find the corresponding moving image, *M*. Since, we already have a set of candidate atlases from the previous step, we select the best matching atlas from this set as the moving image, *M*, for the registration procedure. In other words, we must decide which cluster 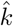 the image to be segmented belongs to, and assign the atlas from that cluster as the moving image. That is to say, the moving image 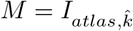 is selected using:

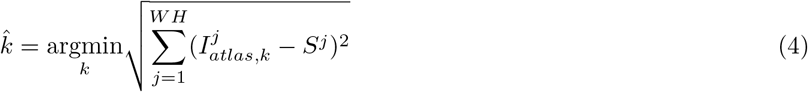

During the registration procedure, the pixels of the moving image are moved as per the accelerated Demon’s equation given by Wang *et al*. [31] in Equation (5). For a moving image, *M*, and a static image, *S*, the additional transformation field, Δ**U**^(*i*)^, for movement of intensity pixels in iteration *i* is calculated as

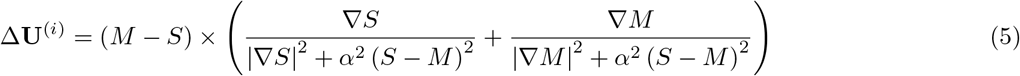

where *α* is the normalization factor that allows the transformation field strength to be adjusted adaptively in each iteration and ∇ computes the gradient for 2D images. Note that the transformation field **U**^(*i*)^ is a vector for each pixel with both an *x* and a *y* component.

The total transformation field after the *i*th iteration, **U**^(*i*+1)^, is the sum of previous transformation field and the additional transformation field calculated in iteration *i* scaled by a value of *s_k_*, speed factor used to speed up the registration process. Equation (6) shows the transformation field update step.
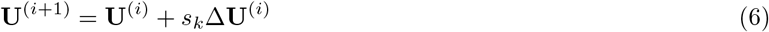

Within each iteration, the total transformation field, **U**^(i)^, is regularized using Gaussian smoothing and the pixels of the moving image are translated with the smoothed transformation. The registration algorithm terminates when we meet the stopping criteria based on correlation of static and moving images given in Equation (7). Here, *ρ*(*S*, *M*)^(*i*)^ is the 2D cross-correlation value of the static and moving images in iteration *i*. We measured the gain in cross-correlation value of static and moving images over ten iterations and stopped the registration process when this gain is smaller than a threshold τ.
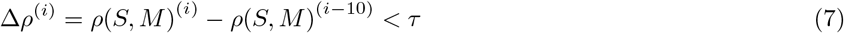

After the convergence of registration process, the segmentation of the moving image is transferred to the static image.

### C. Post-processing

Deformable registration is a pixel-level translation process. This often leads to the formation of holes in the mask. We use morphological cleaning operations, opening and hole-filling, using a disk-shaped structuring element to remove such holes.

## III. RESULTS

We chose mammographic images from the mini-MIAS dataset for evaluation purpose. The dataset consists of 322 images from 161 patients, with left and right mammogram pairs per patient. These mammograms were digitized at 50*μm* pixel edge and downsampled to 200*μm* pixel edge. We downsampled the original 1024 × 1024 images to 512 × 512 for ease of processing.

To standardize the images, we performed a series of preprocessing steps. First, we flipped the left breast mammograms so that all mammograms had the same orientation. Next, images in mini-MIAS dataset are padded on the left and right to make the images square. We removed this padding and translated the the breast region so that it touches the right edge. These two preprocessing steps ensures that the images are in a common coordinate frame.

### A. Evaluation Strategy

We wanted to evaluate how well our method generalizes when there are new images that are not observed during the atlas selection procedure. To do so, we used subject-wise 5-fold cross-validation. In one fold, four partitions of the dataset were used to identify the candidate set of atlas images and the fifth partition was used for segmentation. The subject-wise cross-validation strategy ensures that left and right mammogram images from the same subject were in the same fold. This way, there is no leakage of data between the training and the test set as it ensures that the same subject is *not* in both the training and test set. Indeed, this is to avoid inflated results as could be the case with a simple 5-fold cross-validation strategy [32]. The ground truth segmentation masks for all 322 images were prepared manually and verified by a radiologist. All ground truth masks were prepared using the original 1024 × 1024 resolution images. We used these ground truth segmentation masks for evaluating our final results.

All computations are performed using MATLAB R2014a running on a 64-bit Windows 8 machine with 4 GB RAM and *Intel (R) Core (TM) i5-2430M CPU* @ *2.40GHz* Processor.

**Figure 2:**
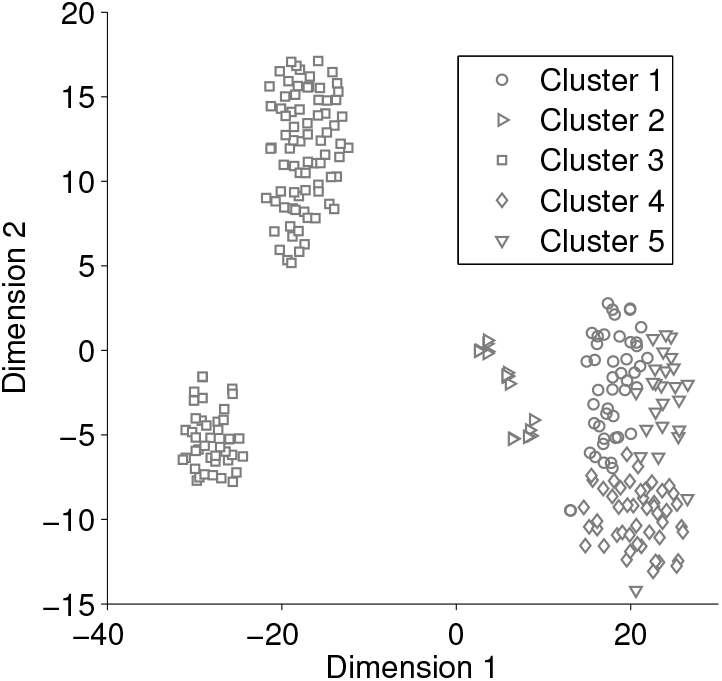
t-SNE projection of mammogram to 2 dimensions. The five clusters found are shown in the projected space.

**Figure 3:**
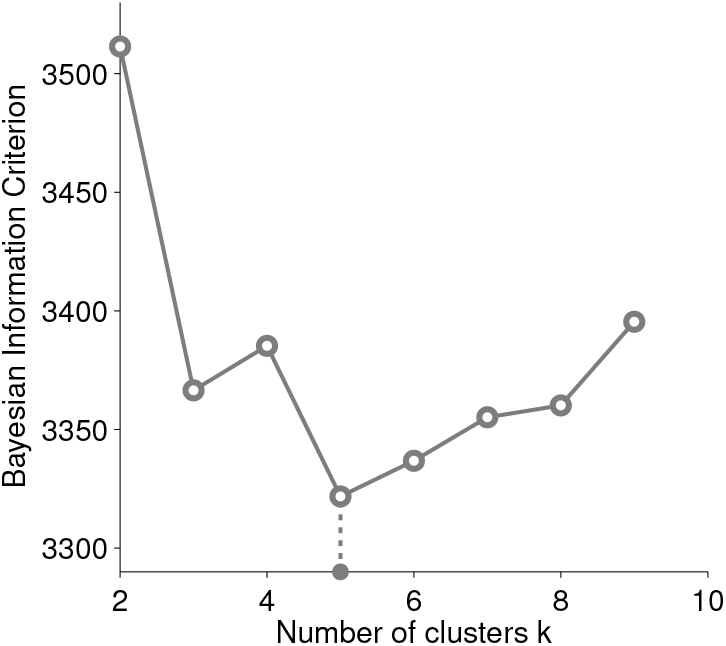
Bayesian Information Criterion (BIC) of GMMs vs. *k*, the number of Gaussians, for the 2D projection data in Figure 2. Minima of the BIC curve suggests presence of *k*\* = 5 clusters.

### B. Atlas Selection

We used t-SNE projection only to identify the number of clusters, *k*\* and not for segmentation. Hence, we used low resolution, 32 × 32, images for this step to reduce unnecessary computational complexity. Here, we present the results from one fold of the cross-validation for illustrative purposes. However, in Section IIIG when comparing against other algorithms, we aggregate the results across cross-validation folds.

Figure 2 shows the t-SNE projection of the mammograms. The corresponding BIC of the GMMs with *k* ∈ [2..9] is shown in Figure 3. The minima of the BIC curve gives the value of *k*^*^ = 5. The 2D projection of the mammograms in Figure 2 shows that the mammograms do in fact belong to 5 different clusters. Figure 4 shows images belonging to the five different clusters. Each row contains images from the same cluster and different rows belong to different clusters. It can be seen that while images belonging to the same cluster are similar in shape and intensity profile, images belonging to different clusters are notably different. It is quite easy to conclude from this figure that performing deformable image registration between images in the same cluster is far easier compared to images from different clusters.

We identified one candidate atlas from each cluster as the image closest to the central cluster image in terms of the Euclidean distance. Figure 5 shows the selected atlas images for this cross-validation fold. The atlas images represent different types of breast images in the dataset. Hence, we argue that data-driven identification of breast shapes and tissue types play an important role in the segmentation process.

**Figure 4:**
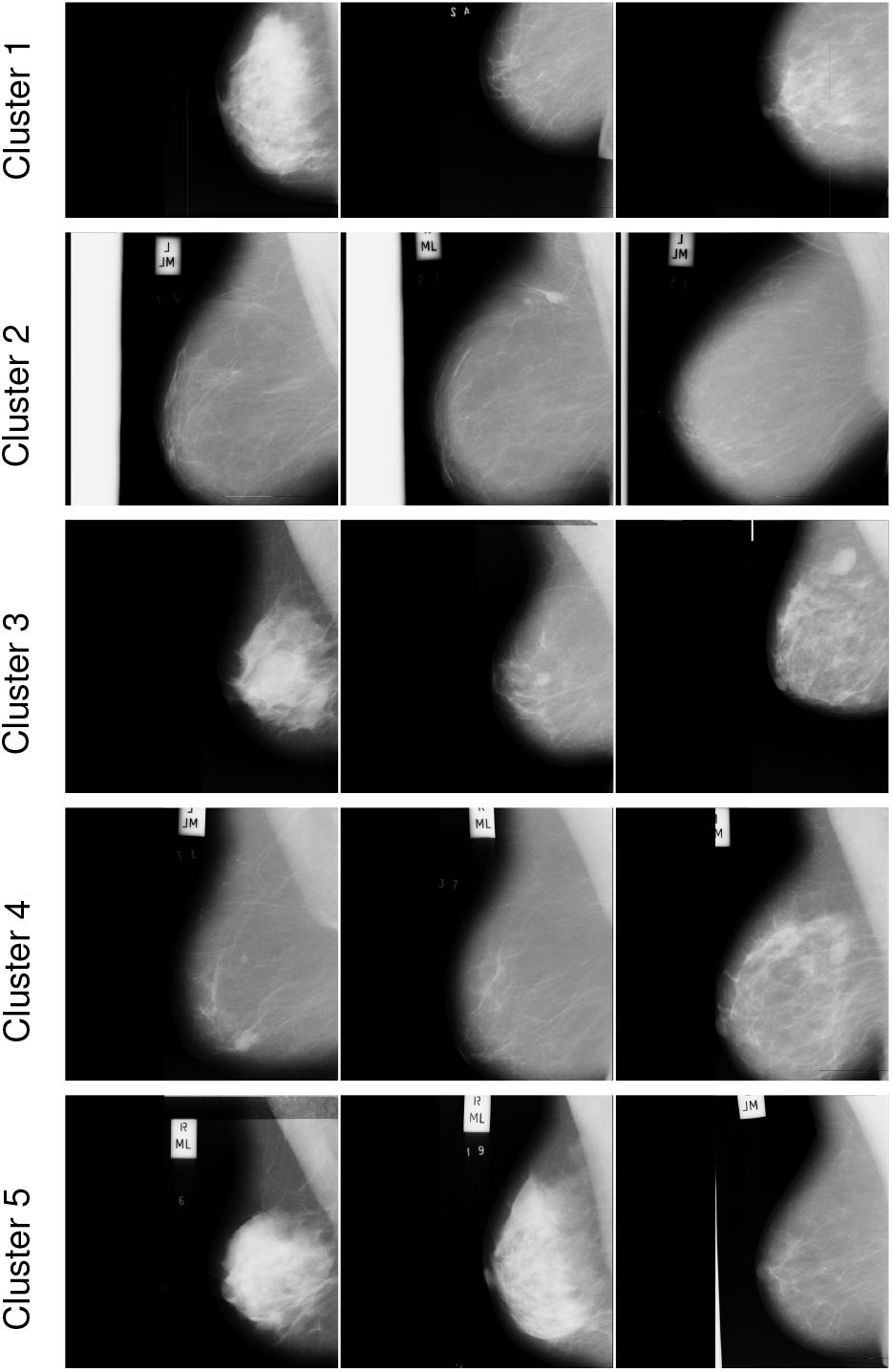
Mammograms belonging to five different clusters of the projection in Figure 2. While images within a cluster are similar, images in different clusters have significantly different intensity profile and shape.

**Figure 5:**
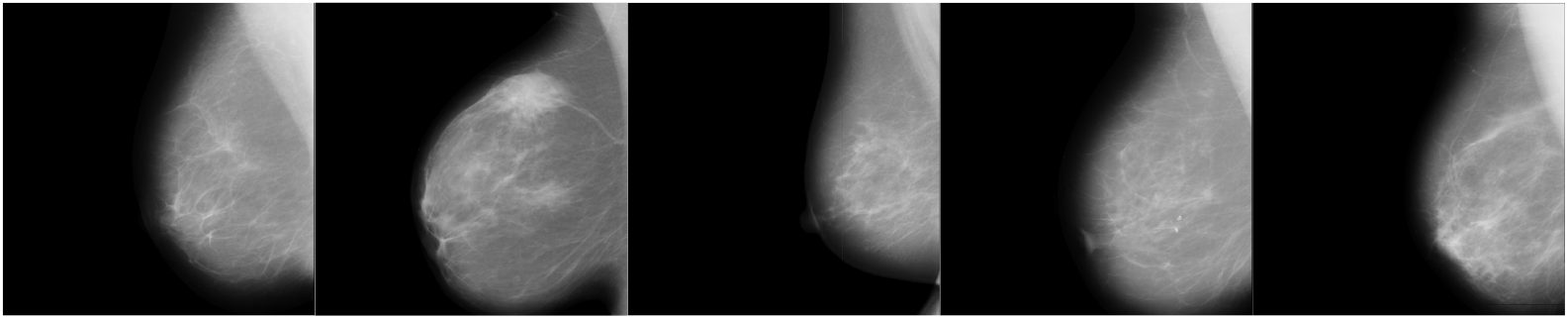
The five atlases selected for each of the clusters in Figure 4. Each atlas represents a different breast shape.

### C. Mammogram Segmentation

We performed the registration using 512 × 512 resolution images. To speed up the registration process, we chose the speedup factor of *s_k_*=4. The average time for the overall registration process was 70.06 seconds per image. The average number of iterations for convergence was 393 per image. The maximum number of allowed iterations was 800.

We used the difference in the 2D correlation coefficient of static and moving images as stopping criteria in Equation (7). We chose the threshold for stopping, *τ* = 0.001. We simplified the equation by choosing the value of normalization constant for transformation field strength *α* = 1. The value of the standard deviation for Gaussian smoothing, *σ*, allows the contours to be curvy, a very small value may result into wiggly contours while a large value may cause the contours to be too smooth. We chose *σ* = 30 as suggested by Jas *et al*. in [23].

**Figure 6:**
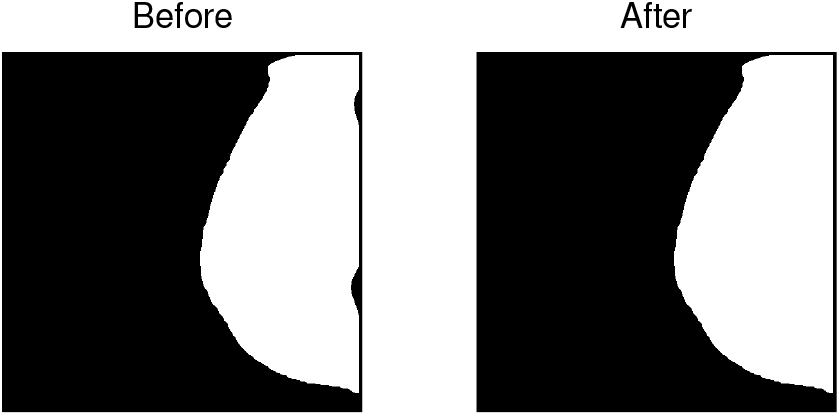
Post-processing step to include breast regions lost during registration process. Here, we show the mask before and after the postprocessing for a particular mammogram.

**Figure 7:**
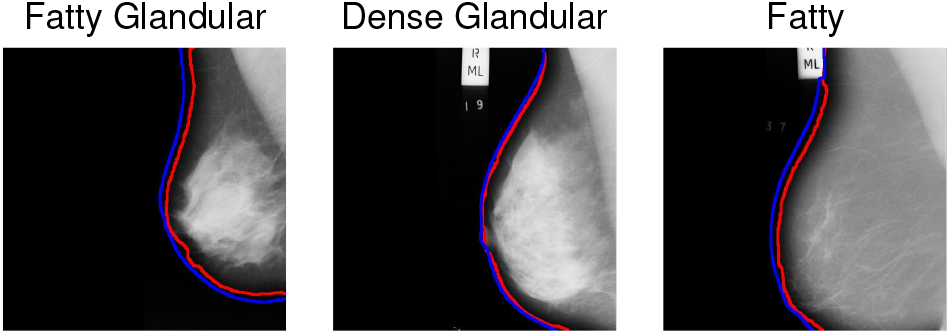
Segmentation results for different tissue types in the mini-MIAS dataset. Each image represents one of the three types of tissues present. Blue boundary shows the ground truth and the red boundary shows the segmentation achieved. It can be seen that the proposed method is able to segment different types of breast tissues fairly close to the ground truth.

### D. Post-processing

Since deformable registration is a pixel-level translation process, it leads to the formation of holes in the mask. We used a disk-shaped structuring element of radius 10 pixels for refining the obtained masks. Also, we observed that some holes were formed along the edges of the image. Recall that in the data preparation stage, we aligned the breast region to the right and top edges. Thus, to recover these regions, we performed a hole-filling morphological operation along the right and top edges. Figure 6 shows a segmentation mask before and after the post-processing step. Finally, for visualization purposes, the segmentation mask was converted into a contour using a Prewitt edge detector and was overlaid on the actual image along with the ground truth.

### E. Segmentation Results for different tissue types

The mini-MIAS dataset classifies breast tissues into three categories: Dense Glandular, Fatty, and Fatty Glandular. Figure 7 shows the segmentation results for different types of tissues present in the mini-MIAS dataset. Each tissue type presents a different shape and intensity profile. Indeed, from visual assessment, we can conclude that the segmentation performs reasonably well irrespective of the tissue type. In Figures 8a and 8b (described in Section III G), we also show that our method outperforms other methods for different tissue types. This can be partly attributed to the fact that the atlases learned in our method represent the variation in the breast shapes across tissue types.

### F. Evaluation Metrics

For evaluating the results, we used Jaccard Index [33] and Hausdorff Distance [34] as the accuracy metrics. Jaccard Index measures the similarity between two sets, *A* and *B*, as the ratio of the number of elements in the set intersection over the number of elements in the set union given by Equation (8).
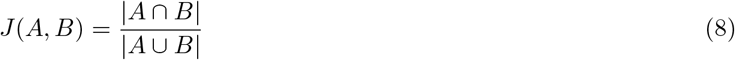

Here, |·| gives the number of elements in the set, also known as the cardinality of the set. In the current context, the set of foreground pixels in the the computed segmentation masks and ground truth masks are A and B respectively. Finally, cardinality of the set is defined as the number of logical ones present in the mask. A higher value of Jaccard index indicates good segmentation while lower values indicate poor segmentation. A value of 1 would indicate perfect segmentation.

**Figure 8:**
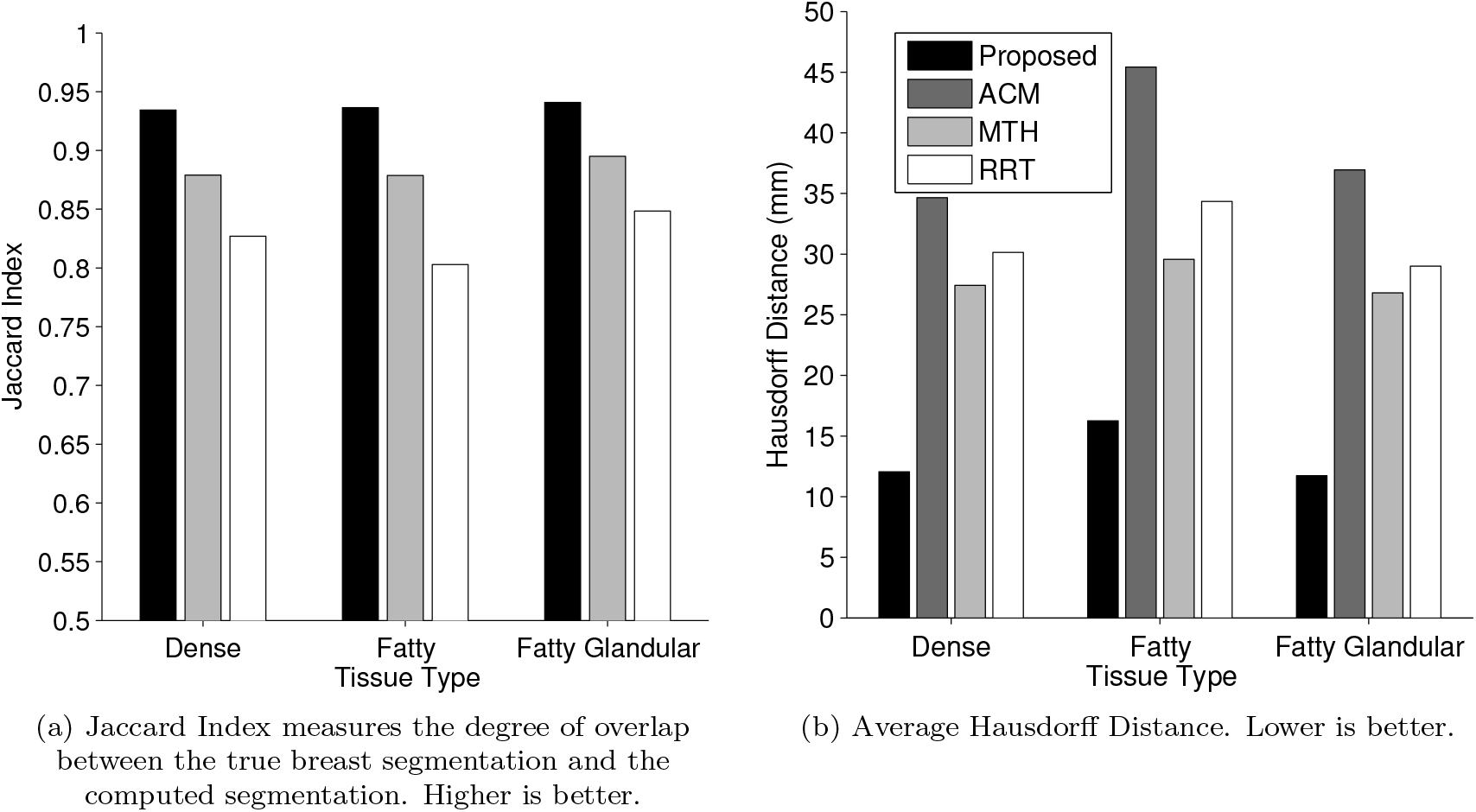
Segmentation results computed on two different metrics for different tissue types in the mini-MIAS dataset. For ACM, we report only the Hausdroff distance as the ground truth masks are not available.

The second metric, Hausdorff distance, measures the maximum of the distances between points of one set to corresponding nearest point in another set. For *a* ∈ *A* and *b* ∈ *B*, the Hausdorff distance, *h*(*A*, *B*), is given by Equation (9), where *d*(*a*, *b*) is the Euclidean distance between the points a and b.
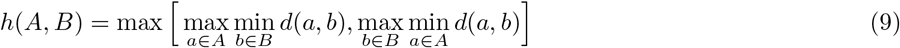

As it measures distances rather than the number of elements, it is affected by the shape of the contours more than the area of the region. Thus, it is a metric complementary to the Jaccard Index. The lower the Hausdroff distance, the better it is. In our analysis, we used the set of coordinates of edge points belonging to the segmentation mask and ground truth mask as *A* and *B*, respectively. The edge points were detected using the Prewitt edge detection method.

### G. Comparison against other algorithms

We benchmarked our algorithm against three state-of-the-art algorithms - Ferrari *et al*.’s Active Contour Model (ACM) [35], Seth *et al*.’s Multi-level Thresholding (MTH) [2] and Abubaker *et al*.’s Row-by-Row Thresholding (RRT) [36]. Ferrari’s method first approximates a breast boundary using a chain code for binarized images. This chain code is used as an input to an active contour model for identification of true breast boundary. Seth’s method of multi-level thresholding divides the histogram into different classes and binarizes the image at the interface of two clusters. Abubaker’s row-by-row thresholding method identifies different thresholds for each row of the image. Each row is binarized using the threshold identified for that row to give the final segmentation. We generated segmentation masks using these methods.

Figure 8a shows the average Jaccard Index and Figure 8b shows the average Hausdorff Distance for different type of tissues present in mini-MIAS dataset. It can be seen from the figure that the performance of the proposed approach is consistently better than the competing algorithms. For each tissue type, the Jaccard Index for our method is higher compared to other methods, while the Hausdorff Distance is lower.

The quality of segmentation achieved by the proposed method and competing methods can also be visually compared in Figure 9. We have marked the segmentation boundaries achieved by these methods and the segmentation boundary of the ground truth in the same image. The red curve represents the segmentation boundaries obtained by the algorithms while the blue curve represents the ground truth. For a fair assessment, Figure 9 considers four different scenarios: 1. The proposed method outperforms competing methods, 2. All methods perform well, 3. Competing methods outperform the proposed method, and 4. None of the methods perform well. It is worth noting that though in some cases, the proposed method is not able to perform well, yet, the segmentation achieved is close to the actual ground truths. We can also notice from the figure that thresholding-based methods are susceptible even to imperceptible salt and pepper noise near the breast boundary. On the other hand, the contour-based method ACM works better although it can sometimes be affected by contours due to external objects.

**Figure 9:**
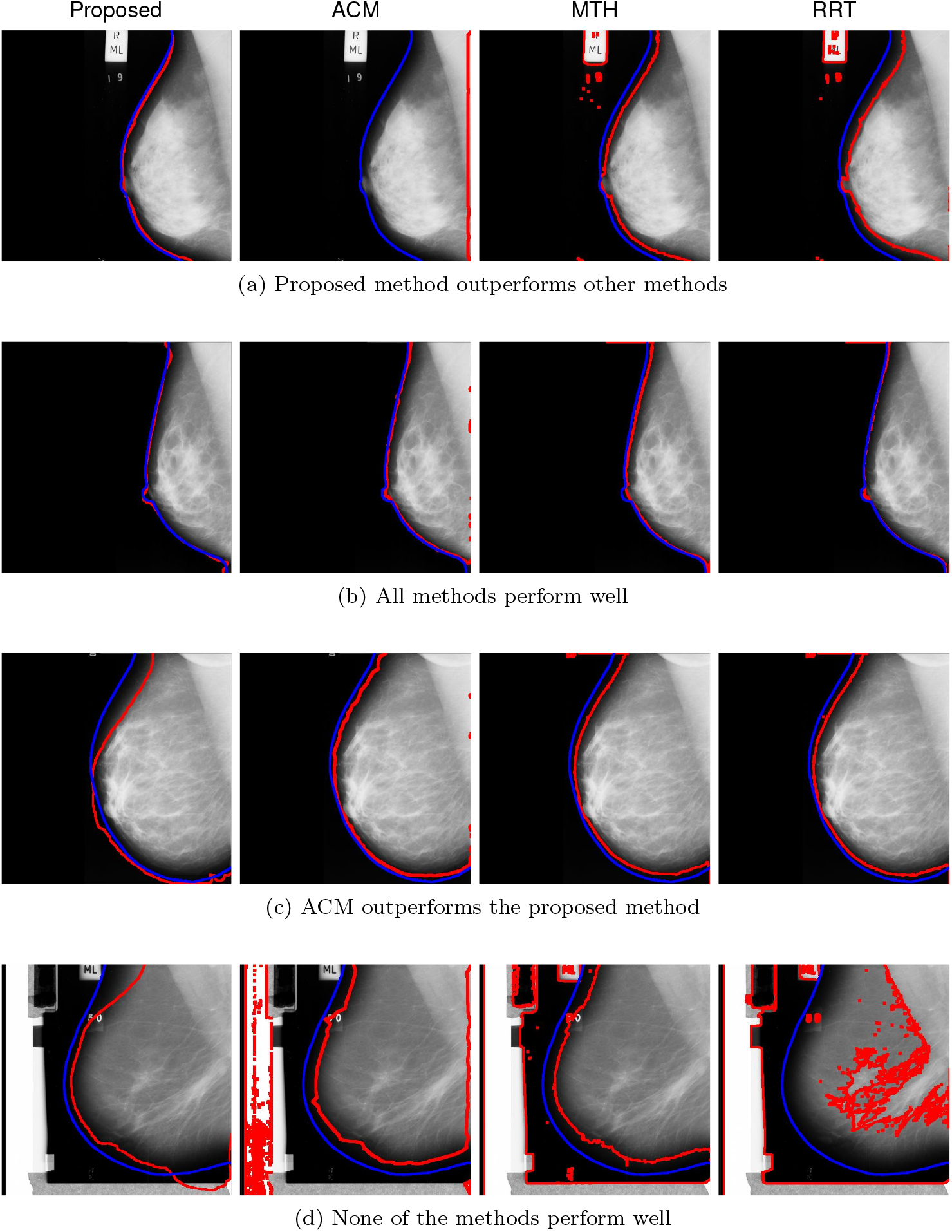
Comparing breast region segmentation obtained using different methods. Methods in the order from left to right are, Proposed method, Active Contour Model (ACM), Multi-level thresholding (MTH) and Row-by-Row Thresholding (RRT). Here the red curve shows the segmentation while the blue curve shows the actual ground truth. We present four performance scenarios here. While each row show a different test image, images within a row show the segmentations obtained using different methods for the same test image.

We compared the performance of these methods quantitatively using the same ground truth masks that were used for visual assessment. For multi-level thresholding and row-by-row thresholding method, the outputs were in the form of masks. However, for adaptive contour model method, the output was the contour of the breast region boundary. Hence, we do not report the Jaccard Index for adaptive contour model. For computing Hausdorff distance, we used the contour for the ACM method and the edges of masks for MTH and RRT. Table I summarizes the evaluated results for different segmentation methods. We can confirm that the proposed method’s performance is certainly better than that of the competing methods in most cases for both the evaluation metrics. Indeed, the average Jaccard Index for the proposed method is the highest while the average Housdorff distance is the lowest. In fact, the proposed method not only performs well on average but also wins against the competing method in a large fraction of images: 79.50% for Jacard Index, and 74.22% for Hausdorff distance.

**Table I:** Comparison of segmentation results obtained with competing methods. Competing methods are Adaptive Contour Modelling (ACM), Multi-level Thresholding (MTH), Row-by-Row Thresholding (RRT)

| Segmentation Method | Avg. Time<br>(s) | Jaccard Index |  |  | Hausdorff Distance |  |  |
| --- | --- | --- | --- | --- | --- | --- | --- |
|  |  | Mean | Std. Deviation | Best Method in<br>% of cases | Mean (mm) | Std. Deviation (mm) | Best Method in<br>% of cases |
| Proposed Method | 70.06 | 0.94 | 0.04 | 79.50 | 13.34 | 12.79 | 74.22 |
| ACM[35] | 18.69 | – | – | – | 38.94 | 14.45 | 3.42 |
| MTH[2] | 0.09 | 0.88 | 0.05 | 8.70 | 27.93 | 14.50 | 15.84 |
| RRT[36] | 0.93 | 0.83 | 0.14 | 11.80 | 31.16 | 15.86 | 6.52 |

## IV. DISCUSSION

We discussed in the Introduction that though there are several approaches to segment breast region in mammograms, due to high diversity in the shape and intensity profile of mammograms (Figure 4), an accurate segmentation is hard to achieve. Most existing algorithms segmented images without considering the variation in image shapes across the database. In this paper, we propose a data-driven approach for segmenting breast region. We have shown in this paper that the proposed method outperforms three state-of-the-art methods. The reason for this is that our method takes into account the shape and profile of breast tissue in mammograms.

We used t-SNE [28] for dimensionality reduction as it preserves clusters during the projection. Specifically, it preserves distances between two points - similar points in the higher dimensional plane remain close in the projected dimensions and dissimilar points remain distant. Empirically, we verified that clusters are indeed preserved in Figure 2. Images sampled randomly from the five clusters resemble each other within a cluster but are dissimilar between two clusters, Figure 4.

One potential limitation of our work is that the optimal number of clusters *k*\* depends on the t-SNE projection. Depending on the initialization of the t-SNE algorithm, the projection would be different and the BIC plot in Figure 3 may yield a slightly different *k*\*. One solution is to use a deterministic algorithm by either fixing the random seed or by using PCA projection for the initialization in 2D. Another alternative is to select the number of clusters *k*\* to be always slightly larger than the estimated minimum. Certainly, more atlas images can only be beneficial for the registration algorithm. This is also confirmed by BIC scores as the BIC for *k* = 4 is much larger than BIC for *k* = 6. Indeed, such heuristics are frequently used in the statistics literature in the context of model selection using cross-validation [37].

Although segmentation based on deep learning is increasingly common [38–41], for it to be feasible on such a task, one would need a large number of segmented training images, even with pre-trained networks. This is not trivially available for mammograms where expert annotations are required. In comparison, our method requires only a small number of segmented mammograms compared to the dataset size, 5–6 images only, *i.e*. approximately 1.5% for 322 image dataset in the present scenario.

Finally, registration is computationally expensive, particularly for high-resolution images that are so common in mammography. However, this is justified due to the high quality of the segmentation obtained. In the present approach, we performed registration on 512 × 512 images. The time taken for the registration process is approximately 70 seconds per image which although greater when compared to other methods, allows to achieve better segmentation accuracy. In the future, we will consider a more efficient registration strategy such as a multi-resolution approach [31] where the deformable registration is performed sequentially from a lower to higher resolution.

## V. CONCLUSION

In this paper, we propose an atlas-based approach for segmenting breast region in mammograms. Mammogram images do not readily lend themselves to segmentation models from computer vision due to being data-limited. At the same time, atlas-based segmentation which is commonly used in medical imaging, cannot be used due to the large variety of breast shapes. Yet, we show that this limitation of atlas-based segmentation can be overcome by combining it with unsupervised techniques like clustering. Our approach is unique in the sense that it combines a traditional segmentation approach: atlas-based segmentation with machine learning techniques of clustering and automated model selection to achieve state-of-the-art segmentation results on mammograms. This approach ensures that the atlas images represent the nature of the data and type of tissue profiles present. We also show that the proposed method outperforms three existing methods on two different evaluation metrics.

